# Impact of random 50-base sequences inserted into an intron on splicing in *Saccharomyces cerevisiae*

**DOI:** 10.1101/2023.06.21.545966

**Authors:** Molly Perchlik, Alexander Sasse, Sara Mostafavi, Stanley Fields, Josh T. Cuperus

## Abstract

Intron splicing is a key regulatory step in gene expression in eukaryotes. Three sequence elements required for splicing – 5’ and 3’ splice sites and a branch point – are especially well-characterized in *Saccharomyces cerevisiae*, but our understanding of additional intron features that impact splicing in this organism is incomplete, due largely to its small number of introns. To overcome this limitation, we constructed a library in *S. cerevisiae* of random 50-nucleotide elements (N50) individually inserted into the intron of a reporter gene and quantified canonical splicing and the use of cryptic splice sites by sequencing analysis. More than 70% of approximately 140,000 N50 elements reduced splicing by at least 20% compared to the intron control. N50 features, including higher GC content, presence of GU repeats and stronger predicted secondary structure of its pre-mRNA, correlated with reduced splicing efficiency. A likely basis for the reduced splicing of such a large proportion of variants is the formation of RNA structures that pair N50 bases – such as the GU repeats – with other bases specifically within the reporter pre-mRNA analyzed. However, neither convolutional neural network nor linear models were able to explain more than a small fraction of the variance in splicing efficiency across the library, suggesting that complex non-linear interactions in RNA structures are not accurately captured by RNA structure prediction methods given the limited number of variants. Our results imply that the specific context of a pre-mRNA may determine the bases allowable in an intron to prevent secondary structures that reduce splicing.

## INTRODUCTION

Gene expression must be tightly regulated in order to ensure optimal levels of protein to support cellular function. This regulation includes the control of splicing, whereby introns are removed from pre-mRNA to form functionally mature mRNA (Parenteau *et al*. 2008; Jacob and Smith 2017; Rose 2019). Splicing of pre-mRNA is carried out by the spliceosome, a large complex of five small nuclear RNAs and over 100 proteins, which is highly conserved in eukaryotes (Plaschka *et al*. 2019; Wilkinson *et al*. 2020). A frequently used model to study splicing is the yeast *Saccharomyces cerevisiae* (Meyer and Vilardell 2008). In *S*. *cerevisiae*, introns are defined by three essential and highly conserved consensus sequences: the 5’ splice site (consensus sequence 5’-GUAUGU-3’), 3’ splice site (5’-[C/U]AG-3’) and the branch point (5’- UACUAAC-3’). While the interactions of these sequences with the spliceosome have been extensively studied (Spingola *et al*. 1999; Qin *et al*. 2016; Bai *et al*. 2018), much less is known about the roles that other intron sequences play in the regulation of splicing.

Studies based on native yeast introns have identified a few key intron characteristics in addition to the consensus sites that influence the efficient and accurate removal of introns. Generally, a shorter branch point to 3’ splice site distance as well as the presence of uridines in this region positively influence splicing (Cellini *et al*. 1986; Coolidge *et al*. 1997; Schirman *et al*. 2021). Conversely, intron GC content and the probability of RNA secondary structures are negatively correlated with splicing efficiency (the proportion of total RNA that is correctly spliced), with both features generally exerting the strongest effects around intron-exon junctions (Yofe *et al*. 2014; Zafrir and Tuller 2015; Schirman *et al*. 2021). However, in some yeast genes, specific secondary structures are required for efficient splicing (Howe and Ares 1997; Rogic *et al*. 2008; Gahura *et al*. 2011; Plass *et al*. 2012). Additionally, a handful of sequence motifs have been identified that either enhance or silence splicing of yeast introns, and these are enriched near intron splice sites (Yofe *et al*. 2014). Studies to date, however, have been limited by the small number of intron-containing genes in yeast (approximately 350), which typically contain only a single intron; other eukaryotes such as humans contain more than 200,000 introns and around eight introns per gene (Spingola *et al*. 1999; Roy and Gilbert 2006; Qin *et al*. 2016). A study that tested several thousand variants by using intron sequences from eleven different yeast species and synthetic combinations of these sequences was still limited by the available sequence space from native introns (Schirman *et al*. 2021). However, this limitation can be circumvented by the generation of large libraries of synthetic sequences, an approach employed to identify key regulatory characteristics of other *cis*-regulatory elements in yeast and mammalian cells (Rosenberg *et al*. 2015; Cuperus *et al*. 2017; de Boer *et al*. 2020; Savinov *et al*. 2021).

To expand the intron sequence diversity available to analyze, we generated a library in yeast with a reporter gene that contains one of ∼140,000 intron variants. The library was based on a shortened version of the *S. cerevisiae ACT1* intron sequence, with a region between the 5’ splice site and branch point replaced by a random 50-nucleotide (N50) element. Using targeted RNA sequencing, we found that a large fraction of N50 elements affected splicing efficiency. Several intron characteristics due to the presence of these elements, including GC content and predicted pre-mRNA secondary structure, contribute to the observed variation in splicing. This study points to an important role of intron sequences in addition to the highly conserved consensus elements in maintaining a secondary structure that is permissive for splicing.

## RESULTS

### Generation of a library of introns with random 50 nucleotide elements

We sought to analyze splicing in *S. cerevisiae* by assessing the effect of a large number of random sequence elements inserted into a common intron, an approach that can generate much greater diversity of intron sequences than found in native yeast genes. We constructed a library of over 750,000 introns, each containing a unique 50-nucleotide (N50) element. The intron background for the library was based on the native *S*. *cerevisiae ACT1* intron sequence of 309 nucleotides. We shortened the *ACT1* intron to 115 nucleotides by removing the centermost nucleotides between the 5’ splice site and branch point sequence; this shorter *ACT1* intron served as a control (Fig. 1A). For the library, 50 nucleotides in the region between the 5’ splice site and the branch point sequence of the shortened intron were replaced by synthetic oligos of 50 random bases (Fig. 1A). The five nucleotides just downstream of the 5’ splice site sequence and the five nucleotides just upstream of the branch point sequence were maintained to avoid effects on splicing efficiency due to changes at sites that can influence the binding of the spliceosome. The N50 oligos had a generally uniform distribution of all four nucleotides (Fig. S1A).

**FIGURE 1.**
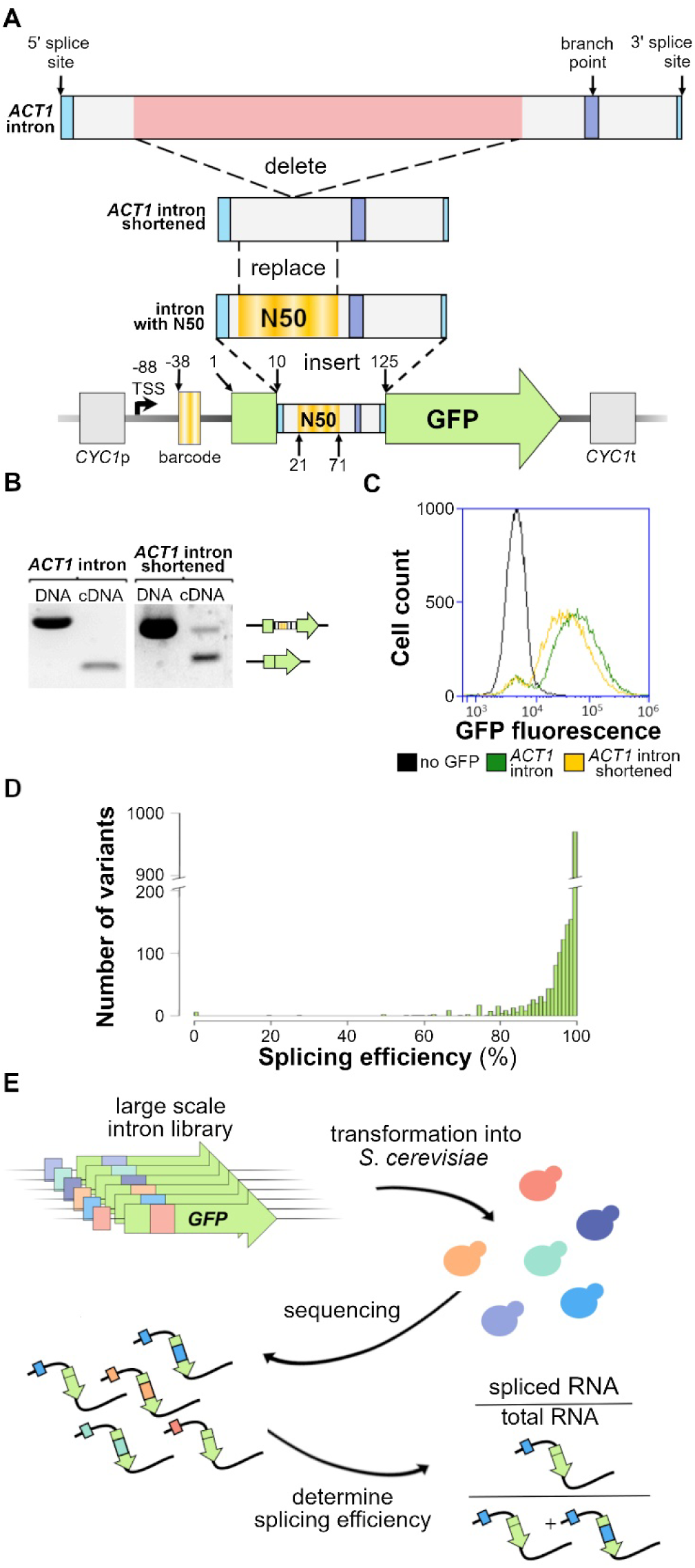
Experimental design and analysis of splicing in controls. (*A*) Design of constructs. The background for the library was created by removing the centermost nucleotides between the 5’ splice site and branch point sequences of the native *S*. *cerevisiae ACT1* intron. A random 50 nucleotide (N50) sequence replaces bases 12 to 61 of the intron between the 5’ splice site and the branch point of the shortened *ACT1* intron. The intron variants were inserted into the GFP gene 10 nucleotides downstream of the start codon with a unique barcode added 18 nucleotides upstream of the GFP coding sequence, which was under control of the *CYC1* promoter and terminator. Numbers indicate the location relative to the GFP open reading frame start site. (*B*) Analysis of splicing of the full-length and shortened ACT1 introns by semi- quantitative RT-PCR using RNA from exponentially growing liquid yeast cultures. PCR using the DNA of the controls are the size references for unspliced transcript (upper band) and spliced transcript (lower band). (*C*) Expression analysis determined by GFP intensity measured via flow cytometry using cells containing plasmids with GFP and the full-length or shortened *ACT1* intron and cells without GFP. (*D*) Splicing of the shortened *ACT1* intron paired with 1905 unique barcode variants as assayed by sequencing. (*E*) Experimental workflow. The N50 intron library was transformed into yeast and RNA was isolated. The splicing efficiency, or the ratio of spliced vs. total transcripts, was measured using targeted sequencing of the intron, and variants were identified by their unique barcode.

Each intron was placed in a Green Fluorescent Protein (GFP) reporter gene at a position 10 nucleotides downstream of the start codon, the same position as the intron in the native *ACT1* gene (Fig. 1A). Transcription of the reporter gene was under the control of the *S. cerevisiae CYC1* promoter and terminator (Fig. 1A). Each intron variant was paired with a unique 15- nucleotide barcode, located in the 5’ untranslated region (UTR), 38 nucleotides upstream of the start codon, such that the barcode remains present in the mRNA after splicing and can be used to identify the pre-mRNA N50 variant for each transcript (Fig. 1A). The constructs were sequenced to match barcodes to N50 elements. The barcodes contained an approximately even distribution of the three nucleotides other than thymine, which was not included to avoid the creation of an upstream start site (Fig. S1A).

To establish the splicing efficiency — defined as the fraction of each reporter gene transcript that was spliced correctly — initially of control constructs, we transformed yeast with plasmids containing either the full-length or the shortened *ACT1* intron within the GFP reporter gene. We analyzed transcript abundance using semi-quantitative reverse transcription PCR. For the full-length *ACT1* intron construct, only the spliced mature transcript was visible by gel electrophoresis (Fig. 1B), comparable to the complete splicing reported for this intron sequence in other gene contexts (Vijayraghavan *et al*. 1986; Agarwal and Ansari 2016). In the shortened intron construct, ∼82% of the transcripts were visible as the spliced form (Fig. 1B). We also assessed protein expression from the GFP reporter using flow cytometry to measure the fluorescence of exponentially growing yeast containing either the full-length or the shortened *ACT1* intron, along with a negative control of yeast with no GFP reporter. Yeast with the shortened intron displayed a reduced fluorescence of 66% relative to yeast with the full-length intron (Fig. 1C), establishing that GFP protein production and RNA splicing efficiency measured by gel electrophoresis were largely concordant for the shortened intron control.

To determine whether the upstream barcode affected splicing, we paired the shortened *ACT1* control with ∼2,000 barcodes and measured splicing efficiency by targeted cDNA synthesis, amplification and sequencing across the 3’ splice junction. On average, the shortened intron was spliced with 96.2% efficiency, with only 0.7% variance between different barcodes (Fig. 1D). Although this efficiency suggests that the barcodes had minimal impact on splicing, 4% of the barcoded controls displayed a splicing efficiency of 80% or less (Fig. 1D). Features of the barcodes may affect splicing, although not GC content of the barcode itself, as no significant differences in splicing were observed among transcripts carrying barcodes of different GC content (Fig. S1B). The analyses of RNA and protein expression demonstrate that the reporter construct is a high-splicing background; however, as it is not spliced as efficiently as with the full-length *ACT1* intron, it can be used to assess insertions that cause either positive or negative effects on splicing efficiency (Fig. 1E).

### Intron variants with random N50 elements display a wide range of splicing efficiencies

We initially characterized the overall splicing efficiency of the intron library. DNA and RNA were extracted from two independent cultures of exponentially growing yeast cells containing the library. To distinguish each transcript, we added unique molecular identifiers (UMIs) to the gene-specific primer used during reverse transcription. Using massively parallel sequencing, we identified each N50 element by its corresponding barcode and determined the abundance of different splicing isoforms by aligning sequencing reads across the 3’ splice junction to either the intron (*i*.*e*. unspliced) or the first exon (*i*.*e*. spliced). The UMIs were deduplicated to remove PCR copies, and unique transcripts were counted as the number of UMIs per variant. To obtain higher confidence data, we included in further analyses only variants that were detected with at least three RNA transcripts and 20 DNA reads (152,514 variants in Rep 1; 193,200 in Rep 2; 141,710 overlap between both replicates (Table S1). The splicing efficiency of N50 variants in the two biological replicates was well correlated (Pearson’s correlation coefficient, R = 0.96; Fig. 2A). Using averages between both replicates in the overlap set, we classified variants into three groups: high (> 80%), moderate (20% to 80%) or low (< 20%) splicing efficiency (Fig. 2A). The library has a large fraction of variants that showed reduced splicing efficiency compared to the control (Fig. 1D and 2A, B). Variants displayed the full range of splicing efficiencies from 0 to 100% (Fig. 2A, B), with 72% of the variants splicing at lower than 80% efficiency (Fig. 2B). This reduced splicing efficiency indicates that features of the N50 elements have a substantial impact on the splicing process.

**FIGURE 2.**
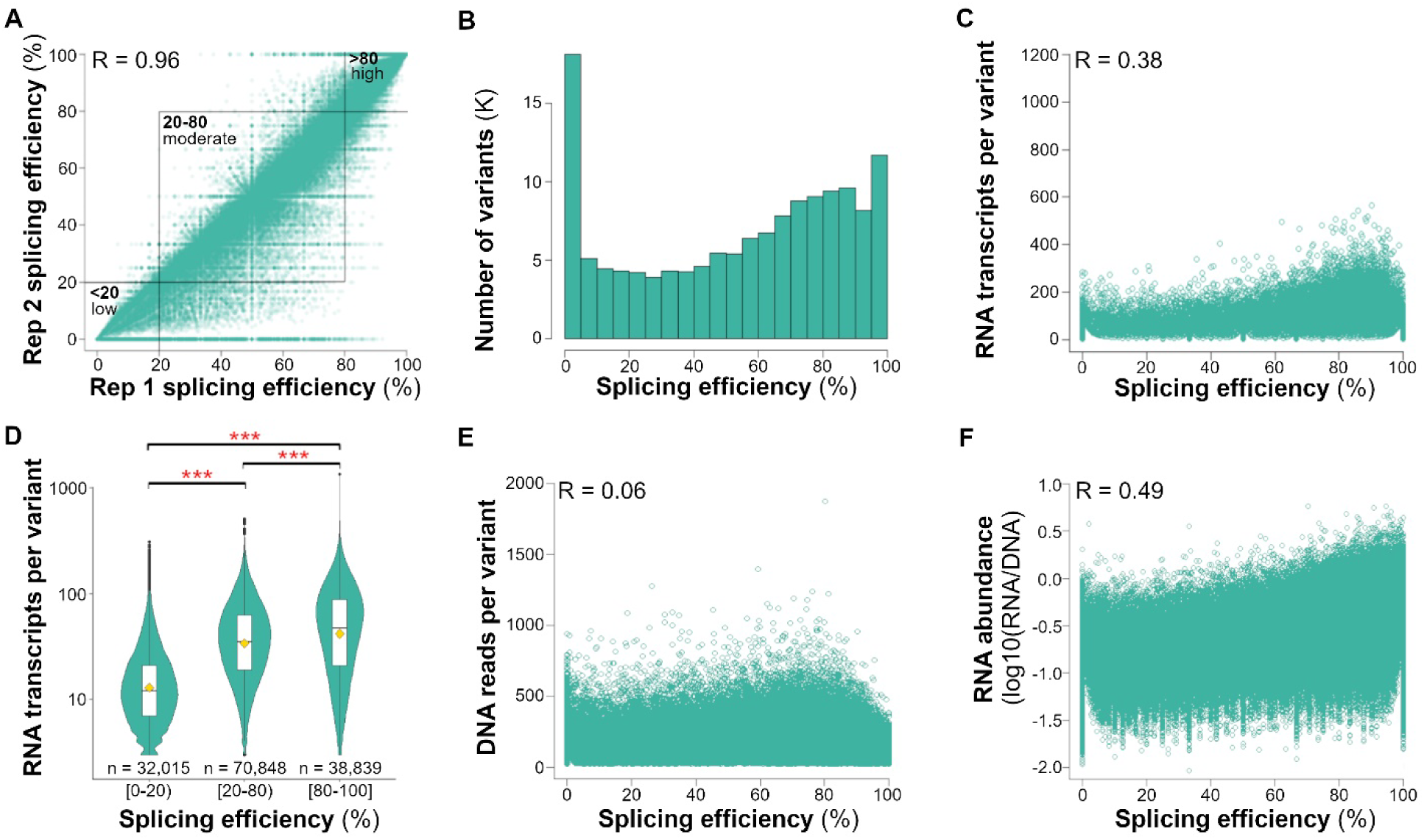
Sequencing analysis of splicing efficiency and transcript abundance in the N50 library. (*A*) Splicing efficiency of 141,710 intron variants present in two independent biological replicates of the intron library. Variants that displayed low (under 20%), moderate (20 to 80%) or high (above 80%) splicing efficiency in both replicates are boxed. (*B*-*F*) Averages of the two replicates are shown without overabundant outliers (> 2,000 reads/transcript, n = 141,702). (*B*) Histogram of RNA splicing efficiency displaying the distribution and range of splicing efficiencies of the N50 library. (*C* and *D*) Total number of RNA transcripts versus splicing efficiency of each variant (*C*) individually and (*D*) averaged across each splicing group (*i*.*e*. low, moderate or high). (*D*) Brackets [ ] indicate inclusive values and parentheses () indicate exclusive values. Horizontal brackets on top indicate significant differences between compared positions using a two-sample t-test with three asterisks denoting p-values < 5^-16^. The gold diamonds indicate the mean for each group. (*E*) Total number of DNA reads versus splicing efficiency of each variant. (*F*) RNA abundance versus splicing efficiency of each variant. RNA abundance was determined by the log_10_ of the total number of transcripts relative to total DNA for each variant. Pearson’s correlation coefficients are denoted by R.

Splicing efficiency and steady-state RNA levels are correlated (Clancy and Hannah 2002; Ding and Elowitz 2019; Schirman *et al*. 2021). We therefore examined the abundance of transcripts containing the random N50 elements. The RNA and DNA levels were moderately correlated (R = 0.57, Fig. S1C). However, the number of RNA transcripts declined as the splicing efficiency of the variants declined (R = 0.38, Fig. 2C). Intron variants that displayed moderate splicing efficiency had on average 25% fewer transcripts, and those with low splicing efficiency a striking 71% fewer, than those with high splicing efficiency (Fig. 2D). Furthermore, unspliced transcripts were 50% less abundant, on average, than spliced transcripts (UMIs per variant, Table S1).

Transcripts with incorrect splicing can be degraded by nonsense-mediated decay (reviewed in He and Jacobson 2015), thereby reducing the accumulation of unspliced transcripts and protecting cells against potentially deleterious proteins produced from these transcripts. The reduced level of unspliced transcripts is also in line with studies that show that inefficient splicing can lead to reduced transcription (Rose and Beliakoff 2000; Rose 2002; Moabbi *et al*. 2012; Agarwal and Ansari 2016). In contrast, the number of DNA reads per variant was negligibly correlated with splicing efficiency (R = 0.06, Fig. 2E). To account for variation in the frequencies of the N50 elements in the population, we normalized each variant’s total RNA to its DNA and termed this “RNA abundance.” The intron library displayed a positive correlation between splicing efficiency and RNA abundance (R = 0.49, Fig. 2F), further supporting the idea that unspliced transcript isoforms have reduced accumulation.

### Intron GC content and secondary structure affect splicing efficiency

To identify sequence features that contribute to the broad range of splicing efficiencies observed in the intron library, we analyzed GC content and the predicted RNA secondary structures across the entire N50 as well as smaller windows of 10 or 15 nucleotides for each N50 element. Generally, introns with higher GC content displayed lower splicing efficiency, with a correlation of R = -0.14 for GC across N50s and R = -0.07 averaged across all 10 nucleotide windows (Fig. S2A, B), similar to prior studies (Galante *et al*. 2004; Wong *et al*. 2013; Schirman *et al*. 2021; Gnan *et al*. 2022). Of the features computed for predicted secondary structures in the library, we specifically analyzed the energy of the structure with the lowest minimum free energy (MFE) (the main structure), and its frequency in the ensemble of all structures. We also examined the free energy and the diversity of the thermodynamic ensemble of structures as well as the base-pairing probability, *i*.*e*. the likelihood of base pairing between each pair of nucleotides, for each library variant. The MFE of the main structure (R = 0.09) and of the ensemble of structures (R = 0.10) were most strongly correlated with splicing efficiency among the predicted secondary structure features using the full N50 sequences (p-value < 2.2 x 10^-16^; Fig. S2C, D). The lower the MFE of the ensemble, *i*.*e*. the more stable the structure of the transcript, the lower the splicing efficiency, suggesting that formation of strong RNA structures influences splicing efficiency. The differences of both GC content and MFE among the three groups of low, moderate and high splicing efficiency were greatest for regions at the edges of the N50 elements relative to the middle (Fig. S2B, C, D). This relationship between strong secondary structures and lower splicing efficiency is similar to results that show that specific secondary structures of the pre-mRNA near or involving the three consensus intron sequences can reduce splicing efficiency and alter splice site selection by hindering spliceosome access to these sites (Halfter and Gallwitz 1988; Goguel and Rosbash 1993; Mougin *et al*. 1996; Singh *et al*. 2007).

The other features of the predicted secondary structures, *i*.*e*. the frequency of the main structure in the ensemble of all predicted structures, the ensemble diversity and the base pair probability, had little to no correlation with the splicing efficiency of the variants (R < |0.02|, Fig. S2E).

### Hexamers enriched in GU reduce splicing efficiency

To identify short sequences in the N50 elements that may affect splicing, we calculated the average splicing efficiency of library members containing every possible six-nucleotide-long sequence. We ranked each hexamer from highest (rank 1) to lowest (rank 4096) splicing efficiency, based on the average splicing of introns containing them, and inspected hexamers more than 2.5 standard deviations away from the mean (Fig. 3A, Table S2). The hexamers with the highest associated splicing efficiency were rich in adenine and uracil (Table S2), in line with the overall negative correlation between GC content and splicing efficiency (Fig. S2A, B).

**FIGURE 3.**
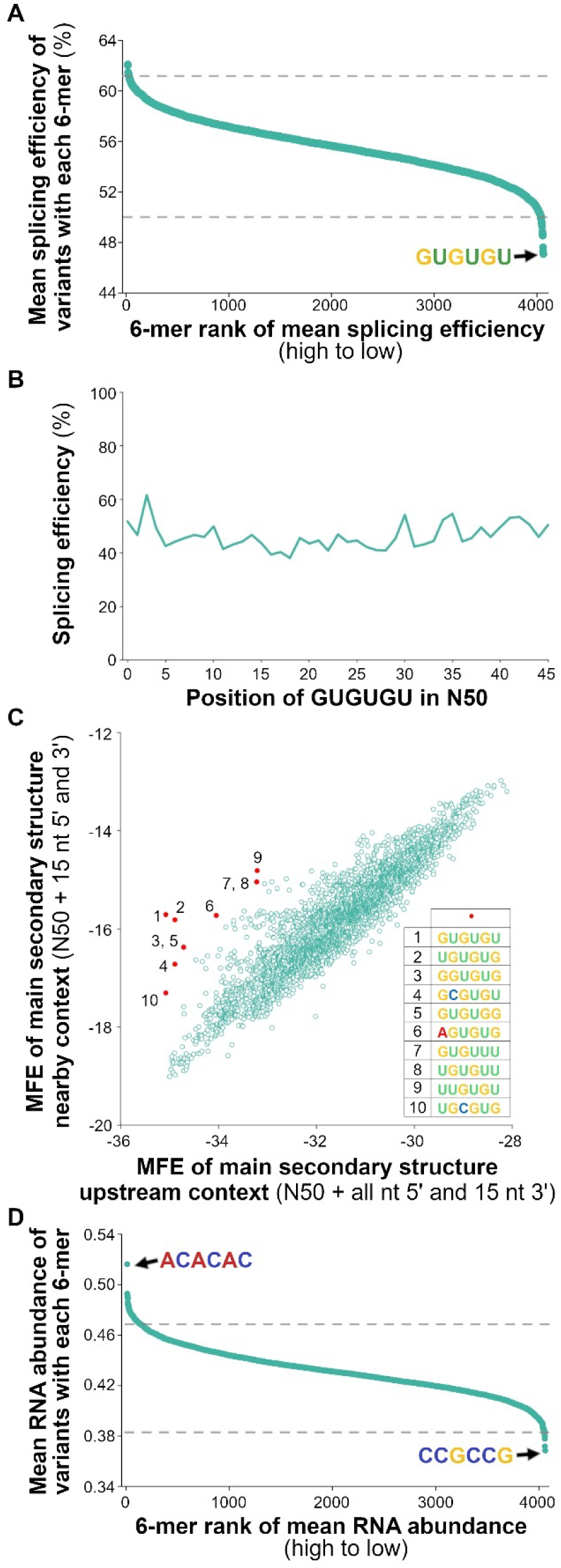
Analysis of all possible hexamers within the N50 library variants. (*A*-*D*) Values averaged between two biological replicates, n = 141,710. (*A*) Average splicing efficiencies of N50s containing each hexamer ranked highest (#1) to lowest (#4096). (*B*) Average splicing efficiencies of N50s containing GUGUGU and starting at each possible position within the N50. (*C*) Average minimum free energy (MFE) of the main predicted structure for the library variants using the N50 sequence and its ‘upstream’ context (entire upstream RNA sequence and 15 bases downstream, 174 bases total) versus the ‘nearby’ context (15 bases each side of N50, 80 bases total). The red numbered dots indicate the specific hexamers listed. (*D*) Average RNA abundance of N50s containing each hexamer ranked highest (#1) to lowest (#4096).

Similarly, hexamers associated with the introns with the lowest splicing efficiency rank were generally GC-rich. However, 20 of the 25 hexamers with the lowest associated average splicing efficiency (ranks 4072 to 4096, 2.5 standard deviations away from the mean) contained at least one GU or UG dinucleotide and 11 contained multiple repeats, with GUGUGU ranked lowest (Fig. 3A, Table S2). The effect of this GU-rich element on splicing efficiency was not dependent on its specific location within the N50, as a similar reduction in efficiency was observed independent of the position of GUGUGU in the N50 (Fig. 3B).

In total, the 25 hexamers with the largest reductions on splicing efficiency were found in approximately 26% of the N50s, due in part to a slight bias for both guanine and thymine in library synthesis (Fig. S1A). We compared the average splicing efficiencies of N50 elements that contain either contiguous GU or UG repeats (*e.g.* GUGU) to those with non-contiguous repeats (*e.g.* GUNNGU). In all cases, N50s with contiguous dinucleotide repeats had on average lower splicing efficiency than those with non-contiguous dinucleotide repeats (Table S2). Although GUGUGU has not been reported to have a drastic effect on splicing when present in native yeast introns, we analyzed the abundance of short sequences in native introns, which might reflect evolutionary selection on sequence content (due to the limited number of native introns, we analyzed pentamers rather than hexamers). GUGUG occurs in native introns more often than 461 other pentamers and UGUGU more often than 778 other pentamers, indicating that GU repeats are not preferentially excluded from *S. cerevisiae* introns (Table S3). Conversely, many pentamers rich in CG or GC dinucleotides are rarely found in native *S. cerevisiae* introns (Table S3). Taken altogether, these results suggest that GC content has a general negative effect on splicing, whereas the effect of the GU repeats is likely due to the specific context of the reporter gene.

We considered whether the GU-rich N50 elements might be forming some secondary structure within the reporter gene pre-mRNA that inhibits splicing. Because of wobble-pairing of guanine with cytosine or uracil, GU-rich sequences could potentially hybridize to a large range of complementary sequences within the pre-mRNA. However, a single CA-rich sequence, ACACACAC, is present in the 5’ untranslated region of the reporter gene, beginning 11 bases upstream of the barcode. To determine if this sequence may be related to the reduced splicing efficiency of introns with GU-rich N50 elements, we calculated the MFE of regions containing the N50 element and variable additional sequence (Table S4, Fig. S3). The correlation of MFE to splicing efficiency was greatest (R = 0.23) using an ‘upstream’ context that spanned from the start of the 5’ UTR through 15 bases downstream of the N50, 174 bases total (Table S4, Fig. S3). Features in this region included the entire 5’ UTR (88 nucleotides), the start of the coding sequence (10 nucleotides), the beginning of the intron (11 nucleotides), the N50 and 15 nucleotides downstream of the N50 (Fig. S3A). Exclusion of the full 5’ UTR sequence using ‘nearby’ (15 bases each side of N50, 80 bases total) or ‘barcode’ (spanning from the start of the barcode through 15 bases 3’ of the N50, 124 bases total) contexts resulted in much lower correlations between MFE and splicing efficiency (Table S4, Fig. S3). Additionally, some hexamers had a much stronger effect on predicted secondary structure when 5’ UTR sequences were included than when they were not (Fig. 3C, Table S2). These outliers included GUGUGU and similar GU-rich sequences, as observed in the lowest ranked hexamers for associated splicing efficiency (Fig. 3C, Table S2). These results support the idea that the GU-rich sequences in the N50 elements form a secondary structure with the 5’ UTR that reduces splicing, suggesting that context is an important consideration in splicing.

We also determined average RNA abundances across N50 elements containing each hexamer and again ranked each hexamer from highest (rank 1) to lowest (rank 4096). N50s containing hexamers with GU repeats were not among those ranked lowest in average RNA abundance, with 29 and 84 sequences ranked lower than GUGUGU and UGUGUG, respectively (Table S2). Rather, the lowest ranked hexamers for RNA abundance were GC-rich, with CCGCCG corresponding to the least abundant RNA on average (Fig. 3D, Table S2).

### Motifs associated with reduced splicing efficiency

We analyzed the N50 elements of RNAs that displayed high, moderate or low splicing efficiency in both biological replicates (Fig. 2A) for sequence motifs that were significantly enriched in each of these three groups. We identified three motifs enriched in the low splicing group, five in the moderate splicing group and one in the high splicing group, when variants were grouped by unique barcodes (Fig. 4A). One of the motifs from the low splicing group and four from the moderate splicing group were similarly found when variants were grouped by unique barcode and N50 (Fig. S4A). To validate these results, we chose representative motifs from each group: (1) UUAGUG, (2) AGGCUUCGGG and (3) GAUGUUUGAAUA from the low-splicing group; (4) UUUGAAUAAUGAC, (5) AAAUGUUCGA, (6) UUUAGAUAUAUGUC, (7) GACUUUAGAU and (8) ACUUAUUU from the moderate-splicing group; and (9) CCGUGUAG from the high-splicing group (Fig. S4A). Validation controls of a splicing enhancer (GUACAUGU, designated Y3) and a silencer motif (UUUGUGUA, designated Y1) identified from native yeast introns as well as a mutated motif that acts as a silencer (UUUAUGCU, designated Y2) were chosen from Yofe *et al*. (2014) (Fig. 4A).

**FIGURE 4.**
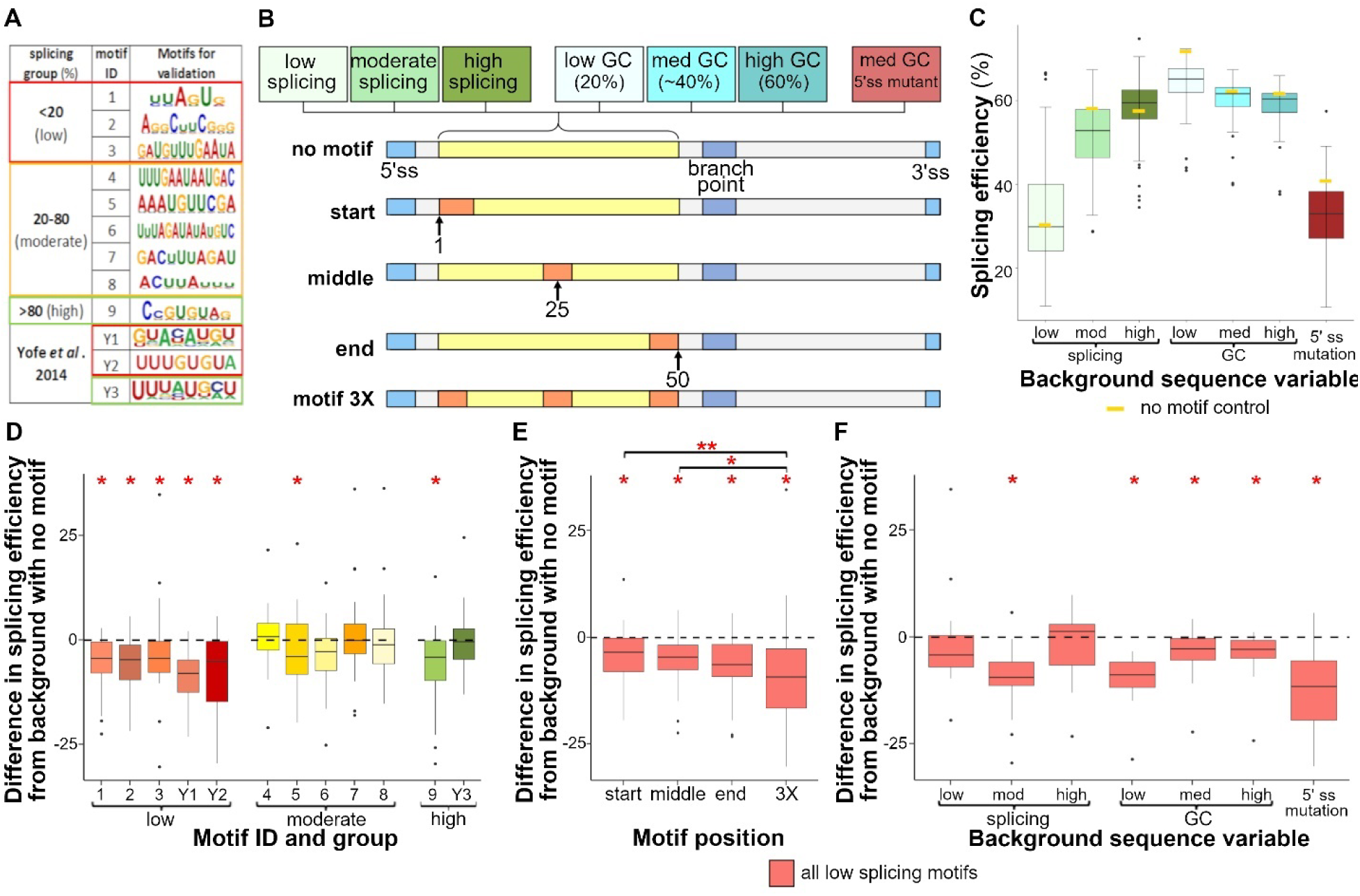
Overview and sequencing analysis of motifs enriched from splicing efficiency status in different intron backgrounds. (*A-F*) Results for the average of the two replicates. (*A*) Motifs that were enriched among the variants with unique barcodes that displayed low (under 20%), moderate (20 to 80%) or high (above 80%) splicing efficiency in both replicates relative to a randomization of those sequences with p-values < 0.05. Also listed are motifs previously reported to reduce (Y1 and Y2) or enhance (Y3) splicing in Yofe *et al*. (2014). (*B*) Overview of the different sequence backgrounds and positions of motifs tested. Numbers indicate the position in the different 50-nucleotide test regions and regions of the introns are denoted ss, splice site; bp, branch point. (*C*) Average splicing efficiency for all motif variants and positions as well as the no motif controls, denoted by a gold rectangle, for each background sequence condition, n = 336. (*D*) The difference in splicing efficiency for each motif variant from the no motif control with the corresponding background sequence, averaged across all positions and backgrounds, n = 28. (*E*) The difference in splicing efficiency for each motif position from no motif control with the corresponding background sequence, averaged across the five low splicing motifs and all backgrounds, n = 35. Brackets indicate significant differences between compared positions using a two-sample t-test with one asterisk denoting p-values < 0.05 and three asterisks denoting a p-value < 0.0005. (*F*) The difference in splicing efficiency for motifs in each background from no motif control with the corresponding background sequence, averaged across the five low splicing motifs and all positions, n = 20. Single asterisks on the top of the graphs indicate significant differences from the no motif control within each group when using a one-sample t-test with p-values < 0.05.

The enriched motifs were introduced into seven different backgrounds and at three locations within the N50 to test their efficacy in a broad set of contexts (Fig. 4B). The seven backgrounds were: one N50 from each of the three splicing efficiency groups that did not contain any of the motifs; the shortened *ACT1* intron sequence with a medium GC content (40%), and low (20%) and high (60%) GC content versions constructed by replacing bases as needed; and the shortened *ACT1* intron paired with a mutation of the 5’ splice site that substantially reduces splicing (Vijayraghavan *et al*. 1986; Fig. 4B). To assess positional dependency, we placed the motifs at one of three locations in the N50: the start (the initial nucleotide of the motif placed at position 1 of the N50), the middle (the central nucleotide of the motif placed at position 25) and the end (the last nucleotide of the motif placed at position 50) (Fig. 4B). We also assessed constructs that had the same motif present at all three positions as well as each background containing no motif (Fig. 4B). In all, we tested 12 motifs in four positional configurations and seven backgrounds, as well as seven no motif control backgrounds, for a total of 343 unique introns.

To test whether the enriched motifs influenced splicing efficiency independent of context, we first compared the splicing efficiency of the seven control backgrounds (no motif). The backgrounds chosen from the low, moderate and high splicing groups displayed the same overall trends in the validation experiment as in the library experiment (Fig. 4C). For backgrounds with higher GC content, there was a corresponding decrease in splicing efficiency (Fig. 4C), again as in the library experiment (Fig. S2A, B). The control background with the 5’ splice site mutation had a splicing efficiency of 52% (Fig. 4C), similar to the effect of this mutation in the wild-type *ACT1* intron (Vijayraghavan *et al*. 1986). To compare the effect of the motifs in these different backgrounds, we calculated the difference in splicing efficiency of each intron compared to the no motif control for each background. A significant decrease in splicing was observed for the three motifs (1, 2 and 3) from the low splicing group, as well as for the two previously reported splicing-reducing motifs Y1 and Y2 (Fig. 4D). No differences in splicing efficiency were detected for motifs 4, 6, 7 or 8 from the moderate splicing group or from the previously reported splicing-enhancing motif Y3 (Fig. 4D). Motif 5 from the moderate splicing group, and motif 9, from the high-splicing group, were highly variable, but showed a significant reduction in splicing across backgrounds (Fig. 4D). These data suggest that only motifs that strongly reduce splicing maintain their effect independent of the background.

We then assessed the position and background specific effects for all low splicing motifs. The average splicing for all low splicing motifs, including the three from the low splicing group and the two from Yofe *et al*. (2014), were observed in all positions and were stronger when the motifs were present in triplicate (Fig. 4E). In analyzing background-specific effects, we found that these five low splicing motifs displayed lower average splicing efficiency in all backgrounds except the low and high splicing backgrounds (Fig. 4F). These results indicate that the motifs were not as effective when present in a background that had a strong enhancing or reducing impact on splicing. In contrast, the motifs further reduced splicing efficiency in the poor splicing background created by the 5’ splice site mutation (Fig. 4F), suggesting that the 5’ splice site and more 3’ regions of the intron impact splicing in different ways. Overall, the results demonstrate that multiple motifs can affect splicing efficiency in context-dependent and context-independent manner.

### N50 variants and cryptic splicing

Cryptic splice sites are non-canonical transcript locations that undergo splicing aberrantly, thereby competing with and reducing the amount of splicing at the canonical site (Nelson and Green 1990; Cunningham *et al*. 1991; Lesser and Guthrie 1993; Rosenberg *et al*. 2015). To examine 5’ cryptic splice sites in the variants, we reexamined sequences that did not align either to the expected junction sequences of canonically spliced or unspliced variants. The unaligned sequences were aligned to the entire unspliced sequence and were designated as cryptically 5’ spliced if the 3’ end of the junction read aligned anywhere upstream of the branch point and not at the canonical site (Fig. 5A). Cryptic 3’ splice sites were identified by junction sequences that aligned combinations of at least six bases of 5’ spliced sequence and at least three bases of 3’ unspliced sequence (Fig. 5A). Approximately 1% of the N50 variants (1,017 variants) had at least one transcript that was cryptically spliced. Most of the 5’ cryptic splices fall within the first 25 nucleotides of the N50 elements (Fig. 5B), likely because there is a minimum length required upstream of the branch point to form a lariat (Wieringa *et al*. 1984; Smith and Nadal-Ginard 1989) and 5’ cryptic splices in the last 25 nucleotides of the N50 would not meet this minimum.

**FIGURE 5.**
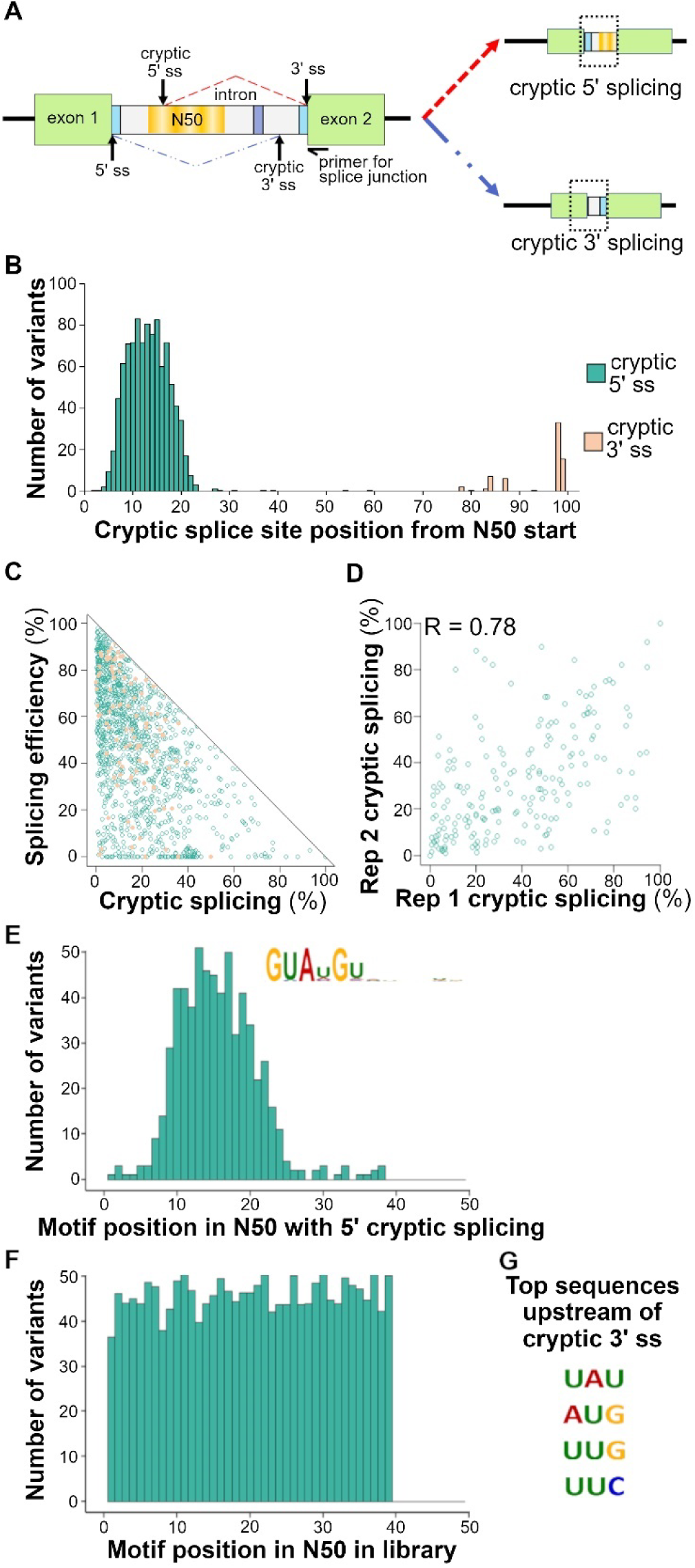
Overview and sequencing analysis of cryptically spliced N50 variants. (*A*) Diagram of the different types of splicing in library variants. Cryptic 5’ and 3’ splicing are depicted by a red dashed line and a blue dash-dot line, respectively. Splicing at a cryptic 5’ or 3’ splice site (ss) results in the inclusion of the 5’ or 3’ end of the intron, correspondingly. The sequencing primer to identify the differentially spliced variants anneals to the junction between the intron and the second exon. (*B* and *C)* Results for the average of both replicates. (*B*) The number of 5’ and 3’ cryptically spliced variants at each position of the N50. (*C*) The splicing efficiency versus the portion of cryptically spliced transcripts in 5’ and 3’ cryptically spliced variants. (*D*) Portion of cryptic splicing per variant found in both independent biological replicates of the intron library (n = 237). Pearson’s correlation coefficient is denoted by R. (*E* and *F*) The number of variants containing the enriched motif at each starting position found in 5’ cryptically spliced variants with (*E*) only 5’ cryptically spliced variants or (*F*) the entire library. (*E*) The significantly enriched sequence motif among the 5’ cryptically spliced variants is listed, p-value < 0.05. (*G*) The four most common sites used for 3’ cryptic splicing among the intron variants.

Cryptic 3’ splice sites were similarly constrained, found only within 30 nucleotides of the canonical 3’ splice site and only at specific locations (Fig. 5B). In the control shortened *ACT1* intron sequence, no 5’ cryptic splicing was observed; however, six of the barcode variants displayed 3’ cryptic splicing (Fig. S4B). The 3’ cryptic splicing in these control sequences occurred at the same location as the top two locations of 3’ cryptic splicing for the library variants, indicating that only a few positions present a viable splice sequence (Fig. 5B and Fig. S4B). No relationship was found between canonical splicing efficiency and the proportion of transcripts that were cryptically spliced (Fig. 5C). For 25% of the N50 elements associated with cryptic splicing, the cryptically spliced transcripts accounted for the majority of the transcripts (Fig. 5C). Only 23% of the 5’ and none of the 3’ cryptically spliced variants were found in both biological replicates. However, the proportion of cryptic splicing for these 23% (237 variants) was well correlated (R = 0.78; Fig. 5D). The low abundance and lack of replicable 3’ cryptic splicing may indicate that the N50 region had comparatively little impact on 3’ splice site selection. For N50 elements in which 5’ cryptic splice sites were identified in both replicates, we searched for enriched sequence motifs. The only motif identified contains the canonical 5’ splice site, GUAUGU, which was found in 73% of the 5’ cryptically spliced variants. This motif is primarily located in the first half of the 5’ cryptically spliced variants (Fig. 5E), similar to where the transcripts are spliced (Fig. 5B). In contrast, this motif is evenly distributed in the canonically-spliced library variants (Fig. 5F). For the 3’ cryptic splice sites, we found that the three bases most commonly used as the cryptic splice site in the library did not match the canonical 3’ splice sequence (5’-[C/U]AG-3’, Fig. 5G). It is possible that cryptic splicing occurs when another site is present that can be bound and cleaved by the spliceosome, even if it is weaker than the canonical splice site.

### Predicting splicing efficiency from intron sequence using multiple models

To determine sequence patterns in the N50 element that influence splicing efficiency, we trained five convolutional neural network (CNN) models with different architectures. For 140,017 sequence variants (the set of overlapped variants with unique barcodes), we trained on 80% of the data, validated on another 10% and tested performance on the remaining 10%. While these models made predictions that are significantly different from random, none of the tested models were able to explain the majority of the variance that was measured in splicing efficiency (max Pearson’s R = 0.26, Table S5). Down-sampling of the number of sequences in the training set suggested that the CNN models require many more training sequences to learn the patterns that are controlling splicing efficiency.

We also applied linear regression, random forest regression and elastic-net models to relate splicing efficiency to 15 known features of each N50 alone or to these features along with k-mers contained in the N50 (see Materials and Methods for feature details; Table 1 and S6). The simple linear regression outperformed the random forest and elastic-net models. The best linear regression model, which included the 15 features as well as the presence of trimers in the N50, performed almost on par with the more complex CNN models, R = 0.24 vs. R = 0.26, respectively (Table 1 and S5). The most important feature in determining splicing efficiency in this linear model was the predicted MFE of the structural ensemble, followed by the GC-content of the N50, the presence of an additional 5’ splice site, the presence of hexamers that were ranked highest in splicing efficiency, the proportion of cryptic splicing and the presence of hexamers that were ranked lowest in splicing efficiency (Table 1). Due to its inherent sparsity, the elastic-net model trained on sequence k-mers within the N50 was also used to determine which sequence patterns have the strongest influence on splicing efficiency. The two most predictive features were the occurrence of GUG elements (predicted to negatively affect splicing efficiency) and the occurrence of AAC elements (predicted to positively affect splicing efficiency). However, structure predictions did not indicate the formation of specific base-pairing between the N50 and other regions of the pre-mRNA, suggesting either that the predictions lack accuracy or that the formation of secondary structure is insufficient to explain the splicing efficiency.

**TABLE 1.**
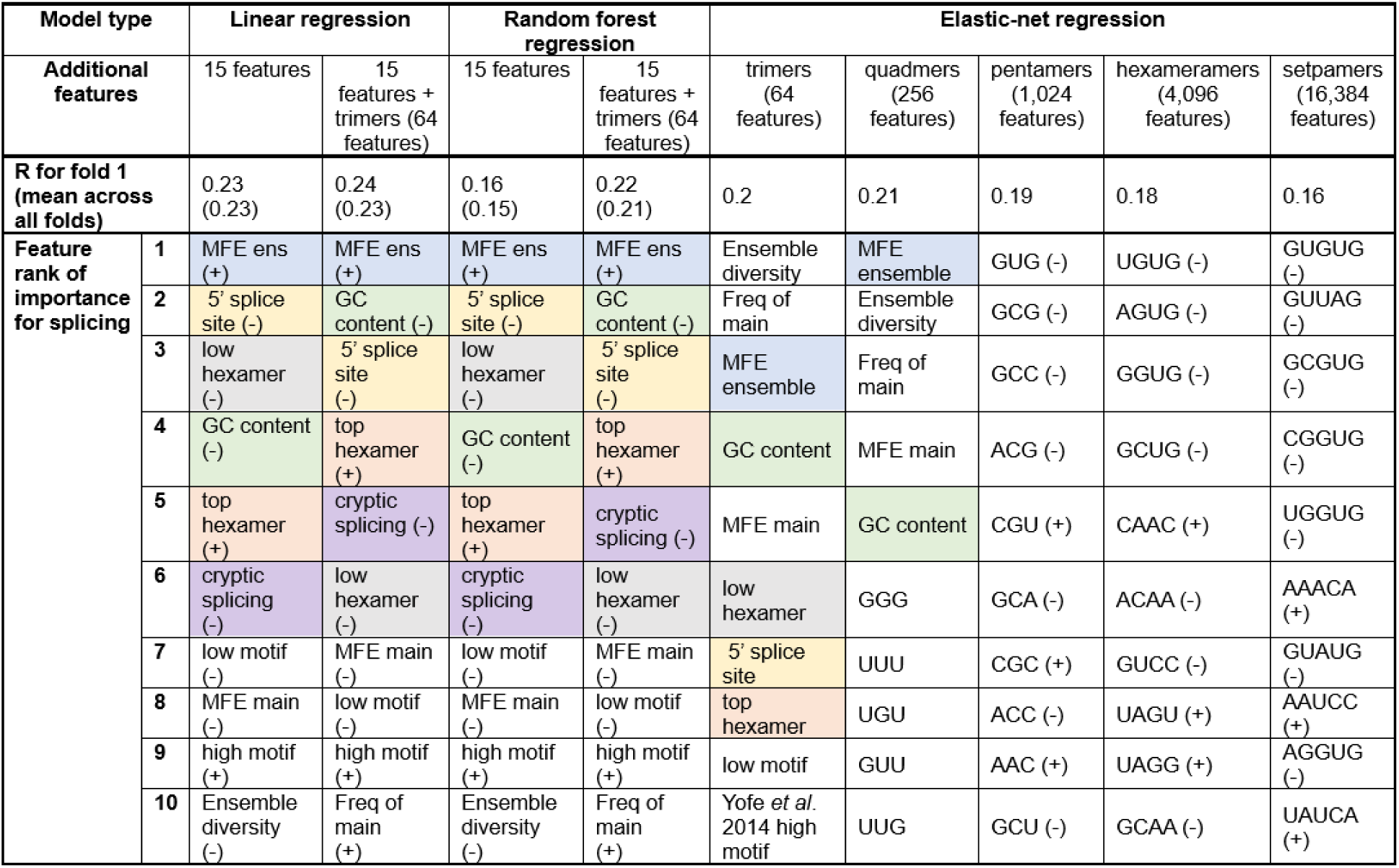
Feature importance and performance of linear models with rank 1 being most important. The top six ranked features in the best performing, *i*.*e*. with the highest Pearson’s correlation (R), linear regression model are colored. The same color denotes the same feature across all models.

## DISCUSSION

This work examined the effect on splicing efficiency in *S. cerevisiae* of approximately 140,000 50-nucleotide-long elements in a shortened *ACT1* intron, with the consensus splice sites and branch point kept intact. The shortened intron control spliced at an efficiency of 80 to 97% of the wild-type *ACT1* intron from which it was derived. The N50 intron variants in this large library displayed the full range of splicing efficiencies, with the majority of variants splicing less efficiently than the control intron. Decreasing splicing efficiency tended to correlate with increasing GC content and stronger predicted secondary structure. These predicted structures correlated much more with splicing efficiency when the 5’ UTR was included in the predictions; in particular, introns whose N50 elements contain hexamers with multiple GU dinucleotides had a dramatic increase in predicted secondary structure when the entire upstream sequence was analyzed in structure calculations, compared to when only 15 nucleotides upstream of the N50 were used. For example, GUGUGU elements went from hexamer rank 2167 to rank 2 (the second strongest predicted minimum free energy, MFE) when the 5’ UTR was included. This change in MFE suggests that the N50 element interacts strongly within the specific sequence environment of its pre-mRNA. The importance of RNA secondary structure for proper mRNA maturation and regulation has been suggested previously. For example, mutation of pre-mRNA to alter or add structural elements can mask splice sites; alternatively, it can promote splicing by organizing splice sites (Halfter and Gallwitz 1988; Goguel *et al*. 1993; Charpentier and Rosbash 1996; Preker and Guthrie 2006; Raker *et al*. 2009). Furthermore, splicing at specific splice sites can increase when heat shock or mutations disrupt secondary structures around these sites (Meyer *et al*. 2011). These reports together with our findings suggest that differences in secondary structure likely impact intron splice site recognition by the spliceosome and thereby influence splicing efficiency.

Our efforts using a variety of models to determine the sequence patterns that cause the N50 effects on splicing were able to explain only a small fraction of the variance. A likely reason is that structure plays a large role in the observed splicing efficiency, but current RNA structure prediction methods have limited accuracy, with maximum expected accuracy at 88% or lower (Mathews *et al*., 2004). While CNN models successfully learn the importance of specific sequence motifs, for example for RNA-binding proteins, they require many more examples to learn complex nonlinear interactions or combinations of features that are represented in RNA structure, where a multitude of sequences in different arrangements can form the same secondary or tertiary structures. It is also unclear whether the structural features that we extracted from RNA structure prediction software fully capture the essential properties that influence splicing.

Intronic splicing regulatory elements are thought to enhance or suppress splicing by forming a secondary structure or recruiting trans-acting splicing factors to the pre-mRNA that interact with the spliceosome, and their function is often location dependent (Wang and Burge 2008; Wang *et al*. 2012; Ohno *et al*. 2018). We identified three sequence motifs – UUAGUG, AGGCUUCGGG and GAUGUUUGAAUA – that were enriched in low splicing variants and that reduced splicing, albeit to differing extents, when present in multiple intron positions and intron sequence backgrounds. Similarly, two previously reported splicing silencer motifs (Yofe *et al*. 2014) were effective at reducing splicing in multiple intron background sequences in our assay. The mechanism by which these sequence elements influence splicing is unclear, although it may involve binding to a splicing factor.

Introns with decreased splicing efficiency tended to have a corresponding reduction in overall RNA transcript abundance. The correlation of splicing efficiency and transcript abundance may be due to greater degradation of unspliced RNA through nonsense-mediated decay (He and Jacobson 2015), as the reporter construct contained an in-frame stop codon in the intron directly upstream of the N50. Less transcription has been observed when introns are removed by the spliceosome at a lower rate (Patel *et al*. 2002), whereas gene expression is enhanced by the presence of efficiently spliced introns near the beginning of a gene (Moabbi *et al*. 2012), and introns at this location may increase transcript initiation (Gallegos and Rose 2015). Highly structured introns have reduced transcript abundance and exhibit lower splicing fidelity when transcribed by RNA polymerase II mutants with altered RNA elongation rates (Aslanzadeh *et al*. 2018).

Some library variants exhibited cryptic splicing, using 5’ splice sites downstream or 3’ splice sites upstream of their respective consensus sites. Much of the 5’ cryptic splicing can be explained by the presence of another site that matches the consensus splice site sequence, specifically within the first half of the N50 region. However, not all variants with a second consensus 5’ splice site sequence in this region displayed cryptic splicing, and several variants that were cryptically spliced did not contain such a site. Thus, additional sequence features may influence the ability of the spliceosome to access or select a splice site.

Overall, this study supports the idea that the secondary structures of pre-mRNAs have evolved to regulate the splicing efficiency of their introns. By beginning with a slightly enfeebled intron, we found that a large majority of random 50-nucleotide insertions further decreased splicing efficiency, often by a substantial amount. This study thus suggests that sequences outside of the canonical splice sites and branch point can have large effects on splicing efficiency.

However, determining the precise mechanisms involved in these effects will require improvements in the ability to accurately predict secondary structure.

## MATERIALS AND METHODS

### Construction of N50-containing intron library and intron controls

For the assembly of constructs, plasmids were linearized using a restriction enzyme digest and then amplified using inverse PCR with KAPA HiFi DNA polymerase HotStart ReadyMix (Kapa Biosystems). The inverse PCR product was treated with DpnI to digest the template DNA. Oligo fragments were amplified with fewer than 10 cycles of PCR using Phusion High-Fidelity DNA Polymerase to minimize PCR error. The DpnI-treated inverse PCR vector and PCR product inserts were column purified using a DNA Clean and Concentrate Kit (Zymo Research). Inserts were added with 2-fold molar excess to 75 ng of the vector, and the two were combined by Gibson assembly (Gibson *et al*. 2009) using NEBuilder HiFi DNA Assembly Master Mix (New England Biolabs Inc.) in 20 µL reactions. Following incubation at 50 °C for 30 minutes, 2 µL of the chilled assembly product was used to transform 50 µL of DH10β *E*. *coli* cells (Invitrogen) by electroporation. Cells were plated in 1:100, 1:1000 and 1:10,000 dilutions on LB agar plates containing 100 µg/mL ampicillin to estimate the number of transformants. The remaining cells were grown overnight in LB media containing 100 µg/mL ampicillin at 37 °C with shaking at 225 rpm, and plasmids were isolated using a PureYield Plasmid Mediprep kit (Promega).

A green fluorescent protein (GFP) reporter gene containing a frameshift (fs) mutation amplified with primers MP1-F and MP2-R (for primer sequences see Table S7) was inserted into a p415- CYC1 plasmid (which contains AmpR and *LEU2* genes for selection, Mumberg *et al*. 1995) linearized by digestion with BamHI and SalI. For the construction of the wild-type and shortened *ACT1* intron control plasmids, the p415-CYC1-GFPfs plasmid was linearized by digestion with XbaI and amplified with primers MP3-F and MP4-R (Table S7), which removed the GFP frameshift mutation and thereby ensured that only correctly cloned plasmids would produce functional GFP. Oligo fragments were obtained from Integrated DNA Technologies that included a section of 5’ untranslated region (UTR) containing a unique 15-nucleotide barcode (without thymine so that additional start sites were not generated) and an intron located 10 base pairs (bp) downstream of the GFP start site. The oligo fragments containing either the *ACT1* intron or a shortened *ACT1* intron with the centermost 194 bases removed between the 5’ splice site and the branch point were amplified with primers MP8-F and MP9-R (Table S7) and inserted into the amplified vector. We added an IDT oligo fragment containing many random 15-nucleotide barcodes amplified with primers MP11-F and MP12-R to the p415-CYC1-GFP plasmid containing the shortened *ACT1* intron that had been linearized by digestion with SpeI and amplified with MP13-F and MP4-R to assess the effect of the barcodes on splicing efficiency of the control (Table S7).

For the construction of the intron library and the validation set, oligos were inserted into a vector amplified using primers F-MP38 and MP4-R from the plasmid containing the shortened *ACT1* intron that was linearized by digestion with XbaI and BamHI-HF (Table S7). The inserts contained the same sequence as the shortened *ACT1* intron except 50 bases between the 5’ splice site and the branch point replaced with random nucleotides (N50) for the intron library or with specific 50-bp sequences for validation and were amplified with primers F-MP47 and R- MP48 (Table S7). The library oligos were obtained as an Ultramer DNA Oligo pool from Integrated DNA Technologies, whereas the validation sequences were obtained from Twist Biosciences.

### Yeast transformation

The BY4741 yeast strain, which is unable to produce leucine, was struck out from a frozen glycerol stock and grown on YPAD plates at 30 °C. Single colonies were used to inoculate liquid YPAD media and grown overnight at 30 °C with shaking at 225 rpm. The cell density of the overnight culture was determined by measuring the absorbance at 600 nm using a spectrophotometer and was diluted to an OD_600_ of 0.1 in YPAD and grown around 4 to 5 h.

Next, the yeast was transformed with plasmid constructs using the high-efficiency yeast transformation protocol of Gietz and Schiestl (2007). Replicate transformations were pooled and cells were plated at 1, 1:10 and 1:100 dilutions on C-Leu plates to estimate the number of transformants. The remaining cells were grown overnight in liquid C-Leu media at 30 °C overnight. Aliquots of the overnight culture were pelleted and resuspended in C-Leu and 25% glycerol and stored at -80 °C. Two biological replicates were grown independently for each yeast strain.

### DNA and RNA extraction, cDNA synthesis and expression analyses

Yeast glycerol stocks were used to inoculate liquid C-Leu media and the culture was then grown overnight at 30 °C with shaking at 225 rpm. The OD_A600_ of the overnight culture was measured using a spectrophotometer and the culture was diluted to an OD_A600_ of 0.1 in YPAD. The yeast was grown at 30 °C and shaken at 225 rpm until the OD_600_ of the culture measured between 0.5 and 0.7, which was around 5 h. Aliquots of the culture were spun down, and RNA and DNA was extracted using 0.75 mL TRIzol Reagent (Invitrogen) for each 0.25 mL of pelleted yeast cells.

The mRNA was purified from total RNA extracts using Dynabeads Oligo (dT)_25_ (Invitrogen) and eluted from the beads. First-strand cDNA synthesis was performed with a gene-specific primer that contained a unique molecular identifier (primer MP56-2, Table S7) and SuperScript IV Reverse Transcriptase (Invitrogen). qPCR was performed on mRNA samples before and after reverse transcription to verify that samples were free from any plasmid DNA. Gel images of qPCR using DNA and cDNA using the wild-type and shortened *ACT1* intron construct amplified with primers F-MP5 and R-MP6 or F-MP10 and R-MP8, respectively (Table S7), were quantitatively analyzed to determine the proportion of spliced and unspliced isoforms of the cDNA by measuring band intensity using ImageJ version 1.53e.

### Preparation of samples for RNA and DNA sequencing

Sequencing samples were amplified by qPCR for 20 cycles or less and between 600 and 800 relative fluorescence units using Q5 High-Fidelity DNA Polymerase 2X Master Mix (New England Biolabs) to minimize replication errors. Each sample set was amplified with primer combinations that added a unique sample index and Illumina’s P5 and P7 adapter sequences to the PCR product (Table S7). PCR products were pooled and column purified using a DNA Clean and Concentrate Kit (Zymo Research) and quantified using a Qubit 3.0 fluorometer.

Sequencing samples were denatured and diluted for sequencing following the standard Illumina protocol and loaded into a high output v2 NextSeq 500/550 reagent cartridge (Illumina). Paired-end sequencing was performed on an Illumina Nextseq 550 sequencer. We first sequenced the intron library DNA to identify the barcode and N50 pairs using custom primers: read1-MP23, read2-MP24, index1-MP22, index2-MP25 (Table S7). Then we sequenced the DNA and cDNA samples using custom primers that read the barcode read1-MP22, the sequence upstream of the canonical 3’ splice site to identify the splice isoform read2-MP32, the unique molecular identifier index1-MP33 and the unique sample index index2-MP34 (Table S7).

### Sequencing analyses and data filtering

For the barcode-N50 sequence assembly, paired-end barcode and N50 reads were assembled using PANDAseq v2.11 and joined using R v4.0.0. Each DNA and cDNA sequencing sample was sorted by their unique sample index using Bcl2Fastq v2.20 allowing no mismatches per index. Barcodes were aligned to the barcodes identified in the sequence assembly and junction reads were aligned to 33 bases of the accurately spliced or unspliced reference sequences using Bowtie 2 v2.4.4. If a junction read did not align, we aligned the sequence to the region starting in the 5’ UTR immediately downstream of the barcode and ending after the branch point to identify any 5’ cryptic splice sites. To examine 3’ cryptic splice sites, we aligned junction sequences to all possible combinations of at least 6 bases of 5’ spliced sequence or at least three bases of 3’ unspliced sequence to filter out any splice site that was part of the canonical splice site. Each barcode-junction-UMI read was connected and UMIs were deduplicated using custom AWK scripts. Data were filtered so that only variants with at least three UMIs and 20 DNA reads were used for analyses to ensure adequate sequencing coverage. Splicing efficiencies were calculated as the percentage of correctly spliced cDNA sequences relative to the total for each variant. RNA abundance was calculated as the total cDNA sequences relative to the total DNA reads per variant. Identification of motif enrichment was determined among variants that displayed low (under 20%), moderate (20 to 80%) or high (above 80%) splicing efficiency in both biological replicates. These sequences were analyzed using STREME relative to the input sequences shuffled for enriched motifs with a size limit of four to 15 nucleotides and a p-value threshold of 0.05 (Bailey 2021). Hexamer ranks were determined by identifying and averaging the values of interest for all N50 sequences containing each hexamer. Correlation statistics and t-tests were used to determine significance using RStudio v2022.07.2 with R v4.2.2 and the rstatix package. Secondary structure estimates were determined using ViennaRNA v2.5.0 (Gruber *et al*. 2008; Lorenz *et al*. 2011), which is based on RNA structure parameters described in Mathews *et al*. (2004).

### Convolutional neural network training and linear modeling

A linear regression employing 15 pre-selected features and an elastic-net regression using sequence k-mers (*i*.*e*. 3-, 4-, 5-, 6- and 7-mers) within the N50 element were implemented with the Python package scikit-learn v1.2.2. (Fabian 2011). Five different convolutional neural network (CNN) architectures with sequence-structure inputs of various lengths were implemented using PyTorch v2.0 (Paszke *et al*. 2019). The models were trained and evaluated using the unique variants that displayed low, moderate or high splicing efficiency in both biological replicates (n = 140,017 variants). All models were trained on 80% of the data using the average splicing efficiency between the two replicates. We tested performance on a held-out 10%, and for elastic-net and CNN models, we optimized hyperparameters on a validation set consisting of the remaining 10% of the data. The sequences in the test and validation set were selected randomly in a way that every efficiency was represented with equal frequency in the training set.

The linear regression model used the following features for each variant: (1) the N50 GC content; (2) the percent transcripts with cryptic splicing; and if the N50 contains any of the (3) bottom or (4) top 10 hexamers ranked by splicing efficiency; the motifs that were enriched in variants with (5) low, (6) moderate or (7) high splicing efficiency; the motifs that were previously reported from native yeast sequences to (8) reduce or (9) enhance splicing efficiency; a canonical (10) 5’ splice site sequence or (11) branch point sequence, as well as (12) the lowest free energy predicted secondary structure; (13) the minimum free energy for the ensemble of predicted secondary structures; (14) the frequency of the main structure in the ensemble; and (15) the diversity of the ensemble using the N50 and its upstream context. After training, we used t-test statistics to rank the regression coefficients by their statistical difference from zero. The linear regression coefficients, *β̂*, of the linear regression model follow a normal distribution with a variance of *σ*^2^(*X^T^ X*)^-1^ (Hastie *et al*. 2009).

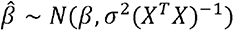

The t-statistic for the coefficients to be non-zero is therefore: 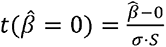, with 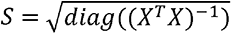, and 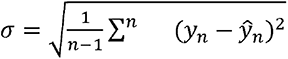, *X* represents the feature matrix of size *(n*, *f*), where *n* is the number of data points, *f* is the number of features, *β* is the true effect, *y* is the measured splicing efficiency and *ŷ* is the predicted splicing efficiency. We performed this ranking for each of the 10 cross-validation folds and determined the order of importance for the features ranking from their average rank across all 10 sets. We also tested a random forest regression model with the same features.

The k-mer elastic-net models used separate sets of k-mers of length 3, 4, 5, 6 and 7 bases. We performed several restarts to determine the optimal regularization parameter alpha (using the validation fold, 10% of the data) for each feature set and reported only the result with the optimal regularization parameter. We also used the t-test statistic to rank the k-mers based on their importance for the predictor.

The CNNs were trained on the one-hot encoding of sequences of different lengths: (a) the 50- base-long random element; (b) 174 bases total, spanning from the start of the 5’ UTR to 15 bases downstream of the N50; and (c) 228 bases total, spanning from the start of the 5’ UTR to 15 bases downstream of the second exon. Although the sequence surrounding the barcode and N50 is constant across the library, the predicted secondary structure can change with the different sequence length depending on the affinity of each N50 element and the barcode to base pair with different regions of the transcript. We also translated the bracket notation of the predicted minimum free energy structure into five structural elements: internal loops, multi- loops, hairpin loops, stacked bases and single-stranded dangling ends using Forgi v2.0 (Thiel *et al*. 2019). These were also added to the one-hot encoding of the three sequence windows as five additional channels. Additionally, the probability for each base in the N50 to be base-paired in the entire 989 base-long transcript was included as an additional input channel to the one-hot encoding.

The CNNs used 200 kernels of length 15 in the first layer, followed by a ReLU activation, and a weighted mean-pooling layer 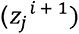 given by the formula:

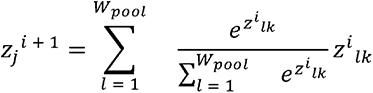

where *i* is the activation from the layer of kernel *k* at position *l* and pooling is performed across the window of size *W_pool_*. The five architectures utilized different numbers of residual convolutional blocks, multi-head attention, additional weighted-pooling layers and fully connected layers to predict the mean splicing efficiency (see Table S5).

## ACKNOWLEDGEMENTS

We thank Tobias Jores for coding and technical advice for sequencing analyses as well as Sayeh Gorjifard for additional coding advice. We thank Christine Queitsch and David Mathews for helpful discussions and suggestions as well as Xinming Tu for the down-sampling analysis of the number of sequences used in the CNN models. This work was supported by grants R01 GM125809, R35 GM139532 and RM1 HG010461 from the National Institutes of Health.

## AVAILABILITY OF DATA AND MATERIALS

The sequencing data are available via the NIH Sequence Read Archive (BioProject ID PRJNA961786). The code for the modeling is available via GitHub (https://github.com/LXsasse/SplicingMPRA).

---

**FIGURE S1.** (*A*) Base frequency in barcodes and N50 sequences of the library constructs sequenced prior to transformation into yeast. (*B* and *C*) Results for the average of the two replicates (n = 141,710). (*B*) Splicing efficiencies of the shortened *ACT1* intron control for specified ranges of barcode GC content as determined by sequencing. Gold diamonds represent the mean for each group. (*C*) Total number of RNA transcripts versus DNA reads per N50 library variant. Pearson’s correlation coefficient is denoted by R.

**FIGURE S2.** (*A*-*D*) Results for the average of the two replicates (n = 141,710). (*A* and *B*) Pearson’s correlation coefficient between splicing efficiency and GC content for all library variants is denoted by R. (*A*) Splicing efficiencies of the intron library variants for specified ranges of GC content across the full length of the N50 as determined by sequencing. Gold diamonds represent the mean for each group. (*B*-*E*) Average values for all library variants that displayed low (under 20%), moderate (20 to 80%) or high (above 80%) splicing efficiency in both replicates. (*B*) The GC content of each 10-nucleotide-long window starting at the 5’ end of the N50. (*C* and *D*) The minimum free energy of the (*C*) main predicted secondary structure in the ensemble and the (*D*) entire structure ensemble for each 15-nucleotide-long window starting at the 5’ end of the N50. (*E*) The base pair probability of each base pairing with any other base in the N50.

**FIGURE S3.** (*A*) Overview of the different sequence contexts used to analyze secondary structure of the RNA transcripts for library variants. (*A*) The sequence used for the ‘nearby’ context spanned from 15 bases upstream to 15 bases downstream of the N50, 80 bases total, the ‘barcode context spanned from the start of the barcode through 15 bases downstream of the N50, 124 bases total, and the ‘upstream’ context spanned from the transcriptional start site (TSS) through 15 bases downstream of the N50, 174 bases total. (*B*-*D*) The minimum free energy (MFE) of the main predicted secondary structure for the (*B*) nearby, (*C*) barcode or (*D*) upstream context versus splicing efficiency of each variant. Pearson’s correlation coefficient is denoted by R.

**FIGURE S4.** (*A*) Motifs that were enriched among the variants with unique barcodes and with unique N50 sequences that displayed low (under 20%), moderate (20 to 80%) or high (above 80%) splicing efficiency in both replicates relative to a randomization of those sequences with p-values < 0.05. (*B*) The number of 5’ and 3’ cryptically spliced barcode variants at each position of the shortened *ACT1* intron.

## TABLES AND TABLE LEGENDS

**TABLE S1.** Overview of the number of unique cDNA and DNA reads sequenced and filtered for the intron library in two independently grown replicates.

**TABLE S2.** The average splicing efficiency (SE), RNA abundance, minimum free energy (MFE) of the main predicted secondary structure for all N50s containing each possible hexamer and averaged for both replicates as well as the number of N50s containing each hexamer and their rank from highest (#1) to lowest (#4096). The sequence used for the ‘upstream’ context spanned from the start of the 5’ UTR through 15 bases downstream of the N50, 174 bases total, and the ‘nearby’ context spanned from 15 bases upstream to 15 bases downstream of the N50, 80 bases total.

**TABLE S3.** Occurrences of each possible pentamer in native *S. cerevisiae* introns.

**TABLE S4.** Pearson’s correlation (R) of splicing efficiency with secondary structure parameters across several regions of the intron library. The sequence used for the ‘nearby’ context spanned from 15 bases upstream to 15 bases downstream of the N50, 80 bases total; the ‘barcode’ context spanned from the start of the barcode through 15 bases downstream of the N50, 124 bases total; and the ‘upstream’ context spanned from the start of the 5’ UTR through 15 bases downstream of the N50, 174 bases total.

**TABLE S5.** Overview of convolutional neural network (CNN) model architectures. Each architecture was individually adapted to the length of the input sequence by adding or removing pooling layers. Each input sequence was used as a sequence in a one-hot encoding (a) or with additional structural channels of sequence plus accessibility (b) or sequence plus five structural elements (c). All models use 200 kernels, convolutional kernel length of 15. The bulleted model parameters appear in the order they were applied to each model. Steps that differ across the three sequence inputs are bold and underlined. Additional terms and abbreviations: the N50 sequence (input sequence #1), a 174 bp sequence from the start of the 5’ UTR through 15 bp 3’ of the N50 (input sequence #2), a 228 bp sequence from the start of the 5’ UTR through the 15 bp 3’ of the intron (input sequence #3) learning rate (Lr), pooling width and stride (W), batch normalization layer (batch norm), dimension of the key, query and value embeddings used in the attention layer (dim. embedding), residual convolutional layer with residual skip connection (residual convolutions), convolutional layer with dilated kernels (dilated convolutions) and width/size of the kernels in the convolutional layer (L).

**TABLE S6.** All data for N50 variants in two biological replicates. Abbreviations: minimum free energy (MFE), lowest energy structure in ensemble (main) of all predicted RNA structures (ens), frequency (freq), div (diversity), replicate 2 (r2), average between both replicates (ave) splicing efficiency (SE), abundance (ab), the bottom (low_hex) or top 10 hexamers (top_hex) ranked by splicing efficiency, the motifs that were enriched in variants with low (low_motif), moderate (mod_motif) or high (high_motif) splicing efficiency, the motifs that were previously reported from native yeast sequences to reduce or (Yof_low) enhance (Yof_high) splicing efficiency, canonical 5’ splice site sequence (fiveP_ss) or branch point sequence (BP), number of RNA transcripts (RNA) and number of DNA transcripts (DNA).

**TABLE S7.** Overview of primers and oligos used for creation, analysis and sequencing of constructs. The letter N represents any base (*i.e*. A, C, G or T) and the letter V represents any base except thymine (*i.e*. A, C or G).

## Notes

### Competing Interest Statement

The authors have declared no competing interest.

